# Conditional Universal Differential Equations Simultaneously Capture Population Level Dynamics and Inter-individual Variation in Human C-peptide Production

**DOI:** 10.1101/2025.01.13.632692

**Authors:** Max de Rooij, Natal A.W. van Riel, Shauna D. O’Donovan

## Abstract

Universal differential equations (UDEs) are an emerging approach in biomedical systems biology, integrating physiology-driven mathematical models with machine learning for data-driven model discovery in areas where knowledge of the underlying physiology is limited. However, current approaches to training UDEs do not directly accommodate heterogeneity in the underlying data. As a data-driven approach, UDEs are also vulnerable to overfitting and consequently cannot sufficiently generalise to heterogeneous populations. We propose a conditional UDE (cUDE) where we assume that the structure and weights of the embedded neural network are common across individuals, and introduce a conditioning parameter that is allowed to vary between individuals. In this way, the cUDE architecture can accommodate inter-individual variation in data while learning a generalisable network representation. We demonstrate the effectiveness of the cUDE as an extension of the UDE framework by training a cUDE model of c-peptide production. We show that our cUDE model can accurately describe postprandial c-peptide levels in individuals with normal glucose tolerance, impaired glucose tolerance, and type 2 diabetes mellitus. Furthermore, we show that the conditional parameter captures relevant inter-individual variation. Subsequently, we use symbolic regression to derive a generalisable analytical expression for c-peptide production.

## 1 Introduction

Many current medical treatments and interventions have been developed and tested in cohorts of individuals, under the assumption that each individual will respond similarly to treatment. However, this one-size-fits-all approach inherently neglects relevant physiological and environmental differences between people, such as genetic variants or acquired exposures that may mediate disease risk or response to treatment. [1]. Furthermore, the observed inter-individual variation may not just be natural variability, but may be indicative of disease progression. Specifically, in the development of type 2 diabetes mellitus (T2DM), the progressive decline of *β*-cell function, responsible for insulin release in response to glucose, is a key characteristic of disease development. [2] Furthermore, residual *β*-cell function is indicative of treatment response and can therefore aid in treatment selection. [3, 4]

In recent years, personalised medicine, where treatments are tailored based on individual characteristics such as genetics [5], or body composition [6], has emerged as a promising approach to improve health outcomes. In particular, in the field of oncology, individual profiling of tumors using machine learning is increasingly being used to select treatment options [7] and has greatly improved the long-term prognosis of patients, demonstrating the potential of precision medicine approaches. [8] However, the direct application of machine learning methods that have yielded success in precision oncology to other medical disciplines has been hampered by smaller sample sizes and a lack of publicly available clinical trial data. [9, 10]

The advantages of these purely data-driven approaches that have been used successfully in precision oncology are that they allow flexible incorporation of various types and sources of data for accurate model output. However, a downside of this flexibility in machine learning models is that the volume of data required to train machine learning models is relatively large. [11] The comparatively small sample sizes collected in human clinical trials have greatly hampered the widespread deployment of machine learning to biomedical problems. [9] In addition, these machine learning models can lack interpretability, particularly in the case of larger neural networks. [12] This interpretability can be retained by using inherently interpretable models, such as ordinary logistic regression, for structured data with meaningful features.

Alternatively, in cases where individual data is limited but biological knowledge is abundant, systems of differential equations are constructed to describe biological processes. These physiologically-based mathematical models (PBMMs) are powerful tools to disentangle the complexity of the physiological basis behind medical measurements. [13, 14] Previous research has demonstrated that the estimation of model parameters in PBMMs from individual measurements can yield an accurate and interpretable explanation of the inter-individual biological variation. [15] While PBMMs are beneficial for studying biological systems, building and validating accurate PBMMs requires a profound understanding of the underlying physiology and can be time-consuming. Consequently, these models typically have a limited scope. [14, 16] Additionally, as PBMMs are constructed manually, unwanted bias can potentially be introduced, especially when complex nonlinear biological behaviours may be approximated with comparatively simple terms.

A promising emerging area of research focuses on the combination of highly plastic machine learning approaches with physiological knowledge in the form of mechanistic models to produce a hybrid model that can be trained with fewer learning examples. In recent years, a number of hybrid approaches have been explored. Physics-informed neural networks (PINNs) [17] are an example, where the loss function of a neural network is supplemented with a set of equations to ensure that the neural network not only fits the data well, but also adheres to known physical laws. In this work, we use the Universal Differential Equations (UDE) framework [18], where the known components of a biological system are described by parameterised differential equations and a neural network is incorporated into the equations to account for unknown components. These UDE models have been shown to be applicable to various biological systems, including the deployment to infer the glucose appearance of a meal [19], as well as a STAT5 dimerisation model. [20] Furthermore, the resulting trained neural networks can be reduced to analytical expressions using a technique called symbolic regression. [18]

Although the application of UDE models has been explored in biomedical contexts, learning the average rate of glucose appearance from a meal, currently conventional training of UDEs cannot directly accommodate inter-individual variability, which is ubiquitous in biomedical data. Although it is possible to train the model on the average data of a population, as has been done in the past, such models are not expected to generalise well to the individual data. Alternatively, it is possible to train a model for each individual separately. However, this approach has some drawbacks. First, limited measurements are often available for separate individuals, making estimation of neural network parameters on individuals highly sensitive to measurement noise, increasing the risk of overfitting. Furthermore, the lack of interpretability of neural network parameters greatly complicates the comparison of trained neural networks between individuals.

In this work, we propose an extension of the UDE framework, labelled conditional UDEs (cUDEs), where trainable person-specific parameters are added as input to the neural network to account for between-subject variability, and the weights of neural network are assumed to be common across the entire population. In this way, variability between subjects is forced into these conditional input parameters, while the neural network parameters learn the global behaviour of the system.

Here, we applied a cUDE model to characterise the insulin production capacity of pancreatic *β*-cells in individuals with normal glucose tolerance, impaired glucose tolerance and T2DM. Our results demonstrate that the conditional universal differential framework derives an accurate representation of the inter-individual variation in c-peptide production. Furthermore, we show that this subject-specific conditioning parameter is strongly correlated with the gold standard hyperglycemic clamp measure of insulin production capacity. We then derived an analytical expression from the conditionally trained network using symbolic regression and showed that the learnt function not only described c-peptide production for people with normal glucose tolerance, impaired glucose tolerance, and T2DM, but also generalised to describe individual c-peptide production in an independent human trial.

## 2 Results

### 2.1 Conventionally trained UDE does not generalise across population

To investigate the ability of a conventionally trained UDE to generalise to meal responses in a population of individuals, a universal differential equation of c-peptide production and kinetics was initially trained on the average meal response. The UDE model is based on a two-compartment ordinary differential equation model describing c-peptide kinetics in the plasma and interstitial space by van Cauter et al. [21]. Here, the van Cauter model was extended by introducing a fully connected neural network to represent c-peptide production in the pancreas (Fig. 1a).

**Fig. 1.**
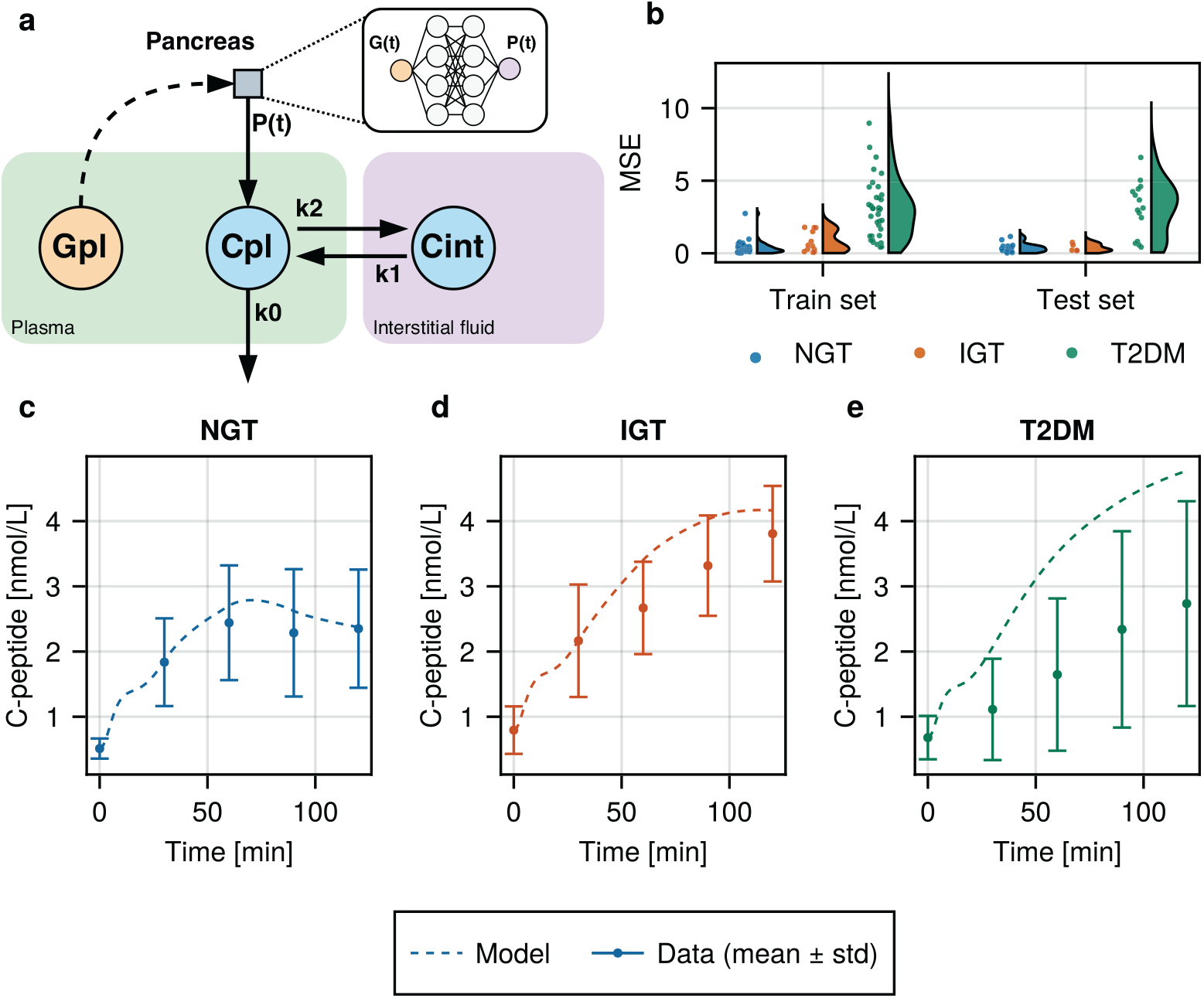
Modeling c-peptide production with a conventionally trained universal differential equation. **a**: Schematic overview of the van Cauter [21] model of c-peptide kinetics, depicting the location of the neural network that describes the production of c-peptide (*P*(*t*)) depending on plasma glucose (*G*^pl^). The blue circles indicate the c-peptide state variables (*C*^pl^ and *C*^int^ for the plasma and interstitial fluid compartments respectively). The orange circle depicts the plasma glucose level. Solid arrows represent fluxes, and dashed arrows indicate stimulation. Each flux arrow is labelled with their respective kinetic parameter. **b**: Mean squared error (MSE) distributions of the UDE model, trained on average response data, on each individual in the used dataset, split by train and test set, and grouped by glucose tolerance status. **c-e**: Mean (circles) and standard deviations (error bars) of the data, and UDE model predictions (solid lines) given the mean data per glucose tolerance condition.

The change in plasma glucose concentration relative to the fasting value at time in provided as input to the neural network *t*, defined as

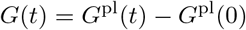

where *G*^pl^(*t*) is the plasma glucose value at time *t*. The output of the neural network is the rate of c-peptide production *P*(*t*).

To train the model, demographic data and plasma glucose and c-peptide trajectories were used from 117 people from a study by Okuno et al. [22]. The data set encompassed three distinct subgroups: people with normal glucose tolerance (NGT), impaired glucose tolerance (IGT), and type 2 diabetes mellitus (T2DM) (S1 Fig). For estimating neural network weights and biases, average c-peptide measurements were used from a training set containing 70% of the individuals. The weights and biases from the neural network trained on the average response were then used in combination with the glucose values and kinetic parameters to predict postprandial c-peptide values. The simulation errors for the individuals in the train and test sets are shown in Fig. 1b, showing comparable performance for the normal glucose tolerance and impaired glucose tolerance groups, but a strong reduction in performance in the type 2 diabetes mellitus group.

Fig. 1c-e shows the resulting model fits, using the average data from each glucose tolerance condition as input. From this figure, we can observe that the model can adequately fit the NGT data but fails to generalise to the IGT and T2DM subgroups. In both the IGT and the T2DM subgroups, the model overestimated the postprandial c-peptide concentrations. This indicates the inability of the single universal differential equation trained on the average response data to account for the progressive decline in *β*-cell function observed in the progression from NGT towards T2DM.

### 2.2 Conditional universal differential equation model of c-peptide kinetics

To be able to capture the inter-individual variability, an additional input parameter was added to the neural network, resulting in a conditional UDE model (cUDE). The resulting neural network embedded in this cUDE model has two inputs. The first inputof the cUDE network is the relative plasma glucose concentration at time t, as in the conventional UDE. The second input is a trainable parameter *β*_*i*_ that accounts for the variability between individuals in the production of c-peptide. The output of the neural network (*P*(*t*)) is the c-peptide production at time *t* (Figure 2a).

**Fig. 2.**
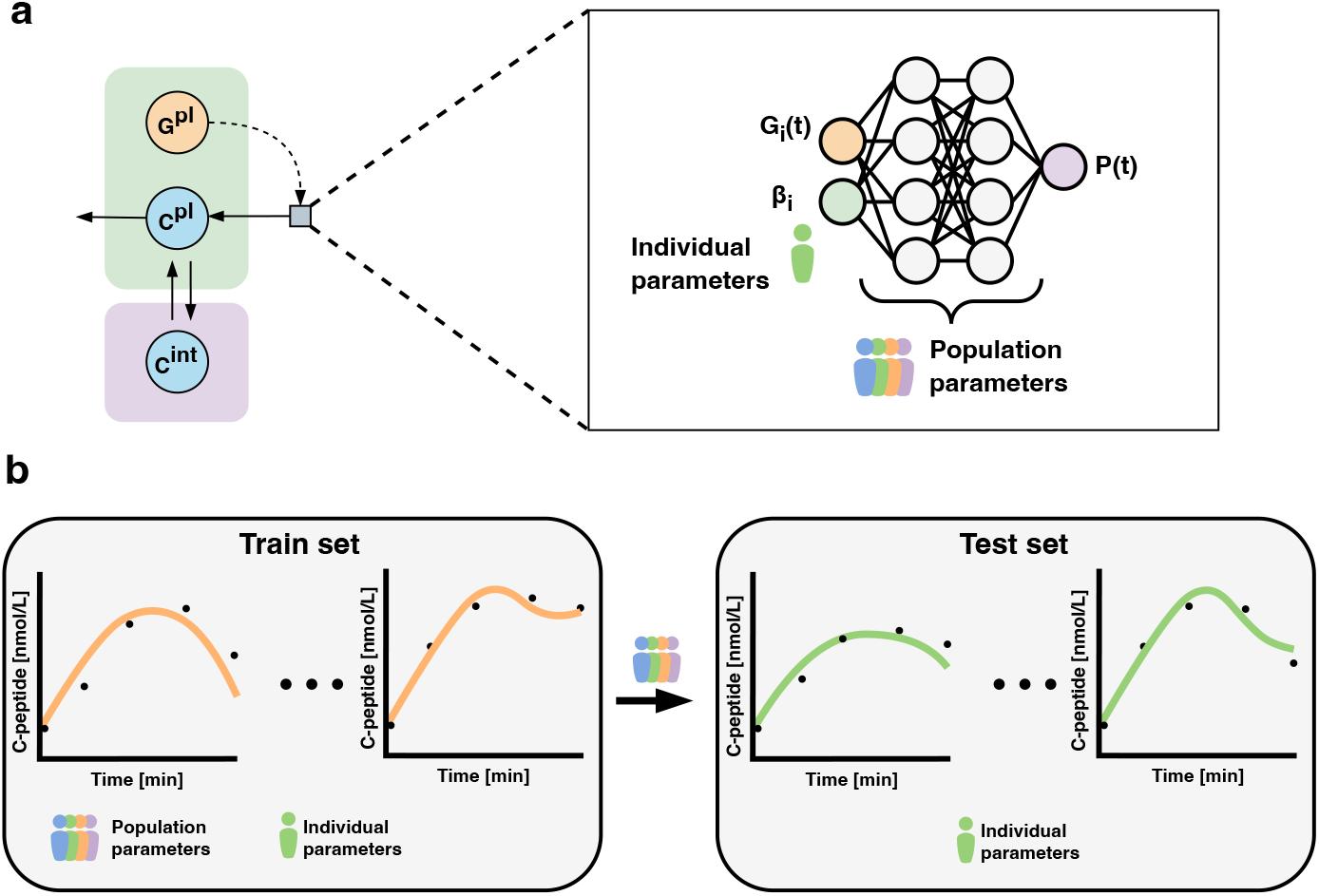
Structure and training procedure of the conditional universal differential equation model used to infer postprandial c-peptide. **a**: Schematic of the conditional UDE model, and the neural network used to estimate personalised c-peptide production. The time-dependent plasma glucose value and a person-specific parameter controlling for inter-individual variability in dose response are inputs to the neural network. The weights and biases of the neural network are estimated population-wide. **b**: Illustration of the training procedure. The dataset is split into a train (70%) and test set (30%). In the train set, both population and individual parameters are estimated. In the test set, population parameters in the neural network are fixed to the trained values and only the person-specific conditioning parameters are estimated from data.

Figure 2b depicts the process of training the conditional UDE. Model selection and training are performed on a subset of 70% the dataset, labelled the ‘train set’. The weights and biases of the neural network for the entire population are trained together with the individual parameters of the train set. The model is then evaluated on a separate test set, where only the individual parameters are estimated, while the neural network parameters are kept constant.

### 2.3 cUDE derives generalisable c-peptide production across population

Fig. 3a-c visualises the cUDE simulation of plasma c-peptide for the individuals in the test set with the median error value for each glucose tolerance condition, showing a good concordance with the measured c-peptide data. This figure demonstrates that the same neural network weights and biases, in combination with a subject-specific conditional parameter, can simulate glucose-driven c-peptide production while accounting for a large part of the inter-individual variability of the c-peptide production. All test fits are shown in supplementary figure S4 Fig. Furthermore, the distribution of model error values across the three groups is shown in 3 d. Compared the to the conventional UDE, the distributions are narrower, especially for the T2DM group. The resulting model fits and error distributions are comparable to the model fits and errors in the train set, which can be found in supplementary figure S3 Fig. In addition, training the cUDE model on various fractions of the train set showed that a train set size of around 29 individuals is already sufficient for training a model, with a comparable mean test error to the current cUDE model (supplementary figure S5 Fig).

**Fig. 3.**
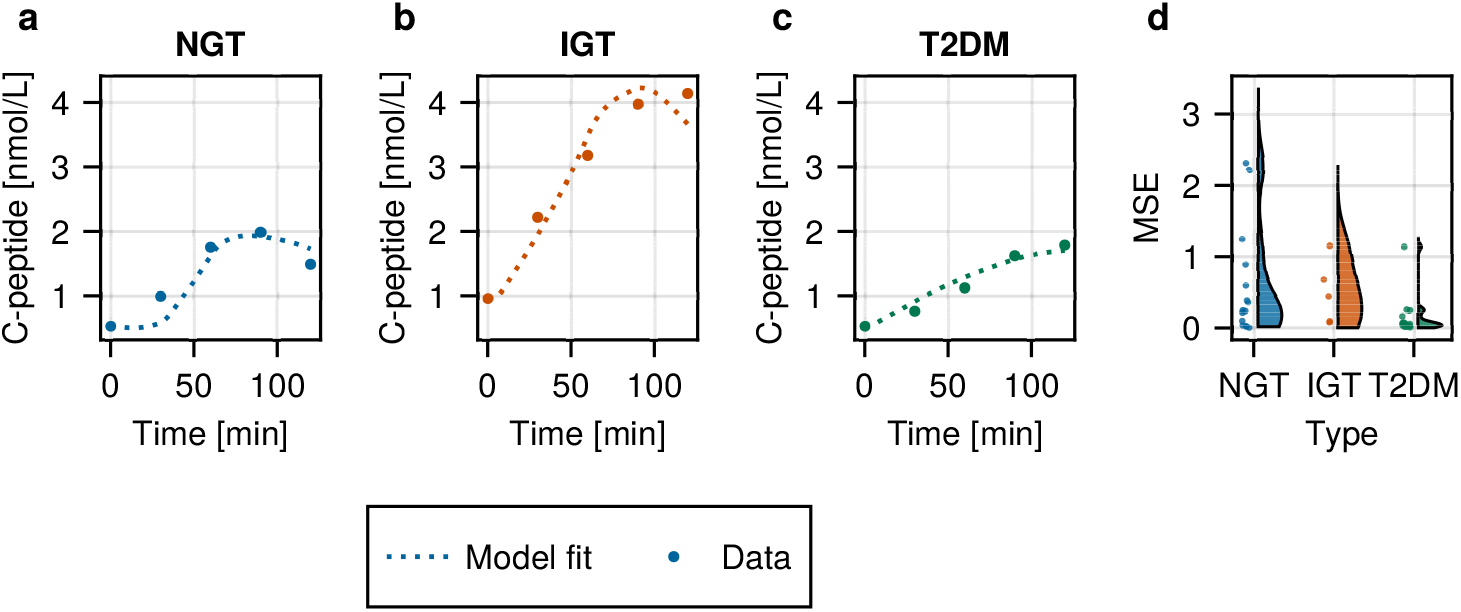
Model fits of the conditional UDE (cUDE) model on the test data. **(a-c)**: Model fit of the individuals with median error value within each glucose tolerance group. Visualisation of all model fits for all individuals in the test set can be found in S4 Fig. Circles indicate the measured c-peptide levels from each individual and dashed lines represent the model fits. **d**: Distribution of mean squared error values for model fits for all subjects in the test subset, separated by glucose tolerance status group.

### 2.4 Conditional training parameter captures inter-individual variation

To investigate the interpretability of the conditional parameter, personalised conditional parameters were compared with subject characteristics, including BMI, age, body weight, and clamp-based measurements of insulin sensitivity and insulin production capacity.

In Fig. 4, the strongest Spearman correlation of approximately 0.81 is observed with the first phase of insulin production measured using the hyperglycemic clamp, the gold standard measure of insulin production (Figure 4 a). A moderate correlation is seen with age (b), while the insulin sensitivity index, measured using a hyperinsulinemic-euglycemic clamp (c), has the lowest correlation of the three.

**Fig. 4.**
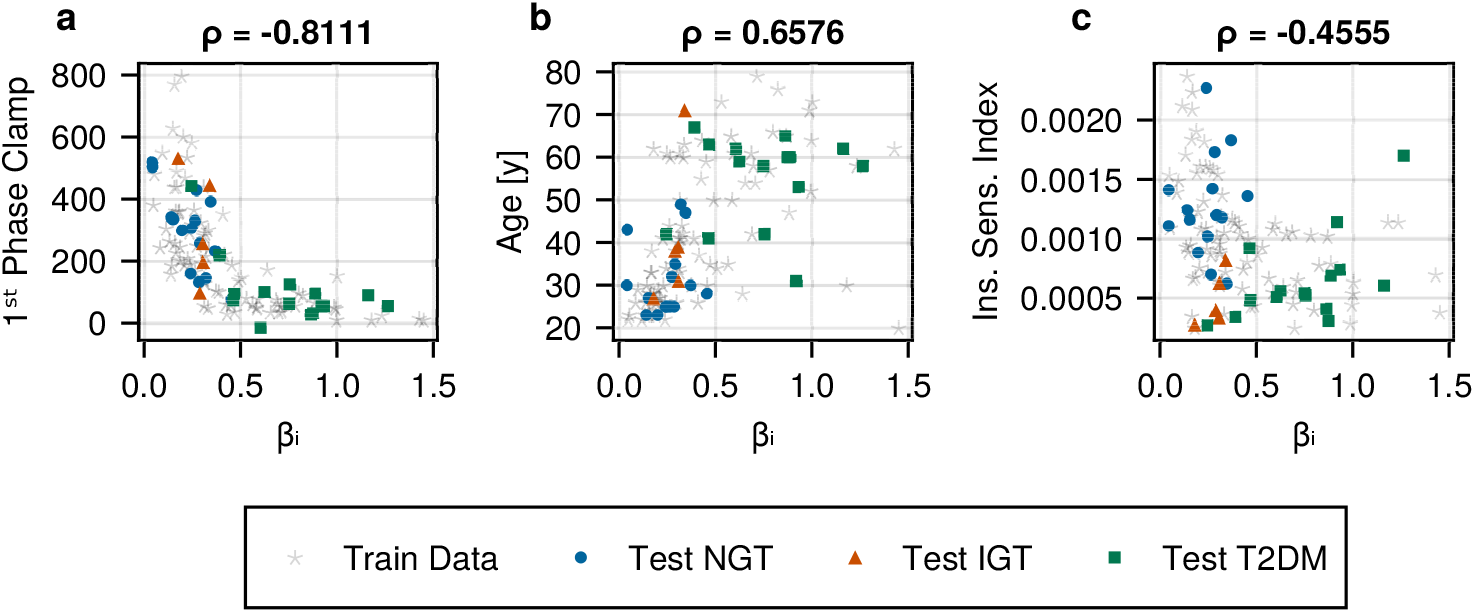
Spearman correlation of conditional parameter *β*_*i*_ with independent phenotypic measurements for the individuals. **(a)** Spearman correlation of the conditional parameter with the first-phase insulin production during a hyperglycemic clamp (S2 Fig) **(b)** Correlation of the conditional parameter with age in years. **(c)** Correlation of the conditional parameter with the insulin sensitivity index measured from a hyperinsulinemic-euglycemic clamp test.

The correlations with body weight and body mass index are low, while the correlations with other measures of insulin production are high, which is shown in the supplementary figure S6 Fig.

### 2.5 Symbolic regression derives a generalisable analytical expression of c-peptide production

As the neural network model remains a black box model, we also sought to replace the neural network with a more interpretable analytical expression. The symbolic regression approach proposed by Cranmer et al. [23] was applied to data sampled from the trained neural network. Subsequently, the derived analytical expression was simplified manually, reducing several fixed constants to a single term (see S7 for a detailed derivation). The resulting expression resembles Michaelis-Menten kinetics and is given as

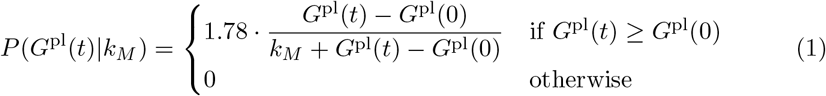

Here, *k*_*M*_ is a trainable parameter, qualitatively equivalent to the *β*_*i*_ parameter learnt by the cUDE. (see S7 for more details on the numerical relation between *β*_*i*_ and *k*_*M*_). The dose-response curves for the neural network and the learnt expression are depicted in supplementary figure S8 Fig.

To evaluate the performance of this learned analytical expression for c-peptide production the neural network of the cUDE was replaced with equation 1. The fully analytical model was then fit to the measured c-peptide data for all individuals by estimating a value for *k*_*M*_.

Fig. 5a-c visualises the model fits of the analytical model model for the individuals corresponding to the median error values per glucose tolerance group. As seen with the cUDE model, the model derived via symbolic regression agrees well with the data across all three groups. The distribution of model fit errors per group (5d) also shows comparable distributions to the model fit errors obtained for the cUDE model. Furthermore, the correlations of the estimated *k*_*M*_ value with insulin production, age, and insulin sensitivity, as shown in figure 5e-g are similar to the previous results obtained with the cUDE model and again display a high correlation with insulin production as measured with the hyperglycaemic clamp.

**Fig. 5.**
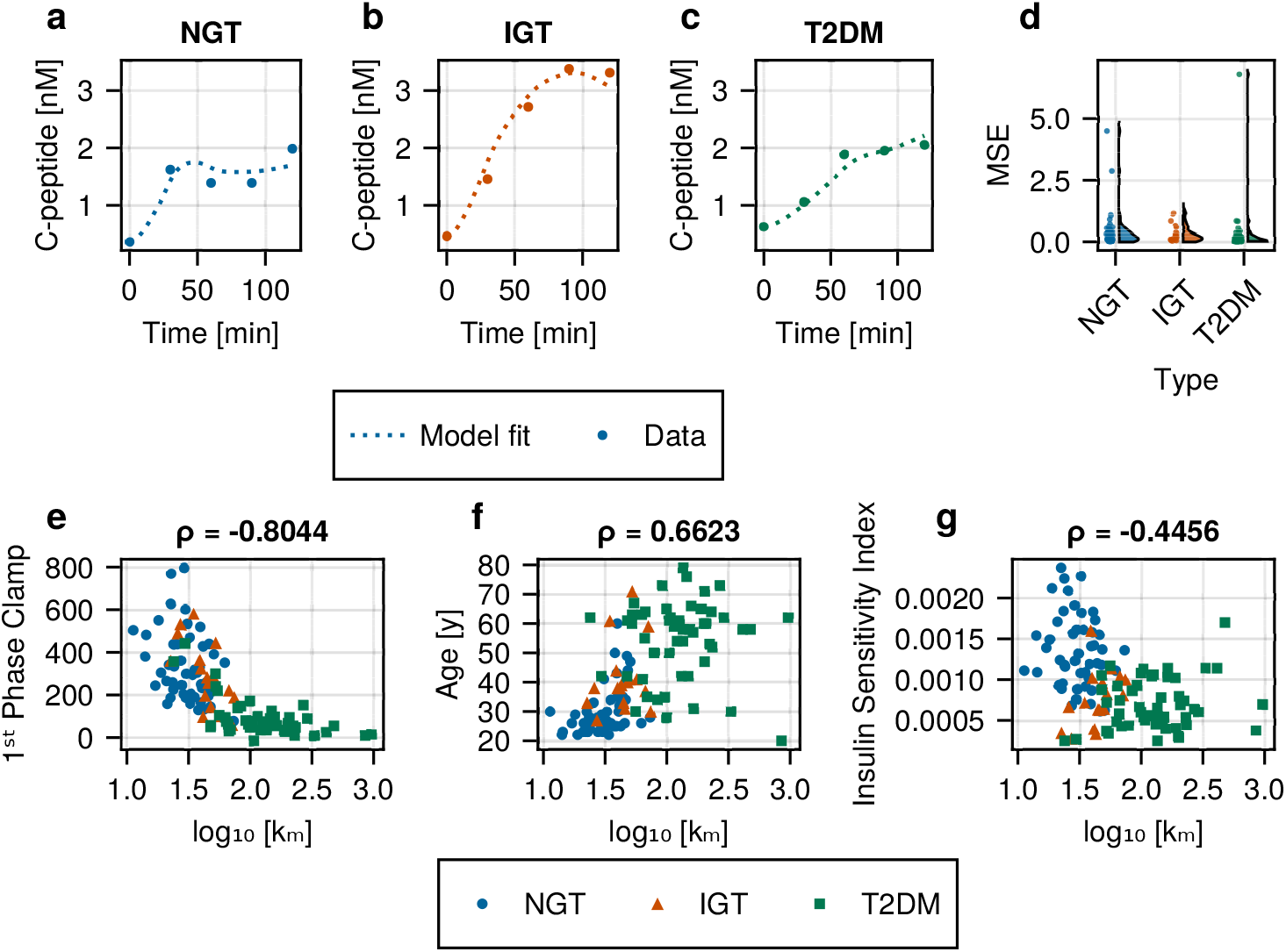
Fit of the analytical model derived using symbolic regression to measured data. **a-c:** Model fit for individuals with the median error value for each glucose tolerance condition. Model fits are shown with the dashed line, measured c-peptide values are indicated with the solid circles. **d**: Model error value distributions split by glucose tolerance condition. **e-g**: Correlation of personalised *k*_*M*_ estimate with **e**: the first-phase insulin production measured during the hyperglycaemic clamp, **f** : age (years), and **g**: the insulin sensitivity index measured using a hyperinsulinemic-euglycemic clamp test.

Finally, to demonstrate generalisability of the model derived from symbolic regression, the analytical model was fitted to glucose and c-peptide measurements collected during an OGTT from a previously unseen dataset.

The model fits for the individuals at the 25^th^, 50^th^ and 75^th^ percentiles of the mean squared error are shown in figure 6a-c respectively. In all three models, the curve shows high concordance with the data. Moreover, despite the original cUDE model being trained for data up to 120 minutes, the learnt analytical term can also reliably simulate plasma concentrations of c-peptide up to 240 minutes postprandially. The distribution of model errors is shown in Figure 6d, indicating that high-quality model fits could be obtained for a large part of the twenty individuals.

**Fig. 6.**
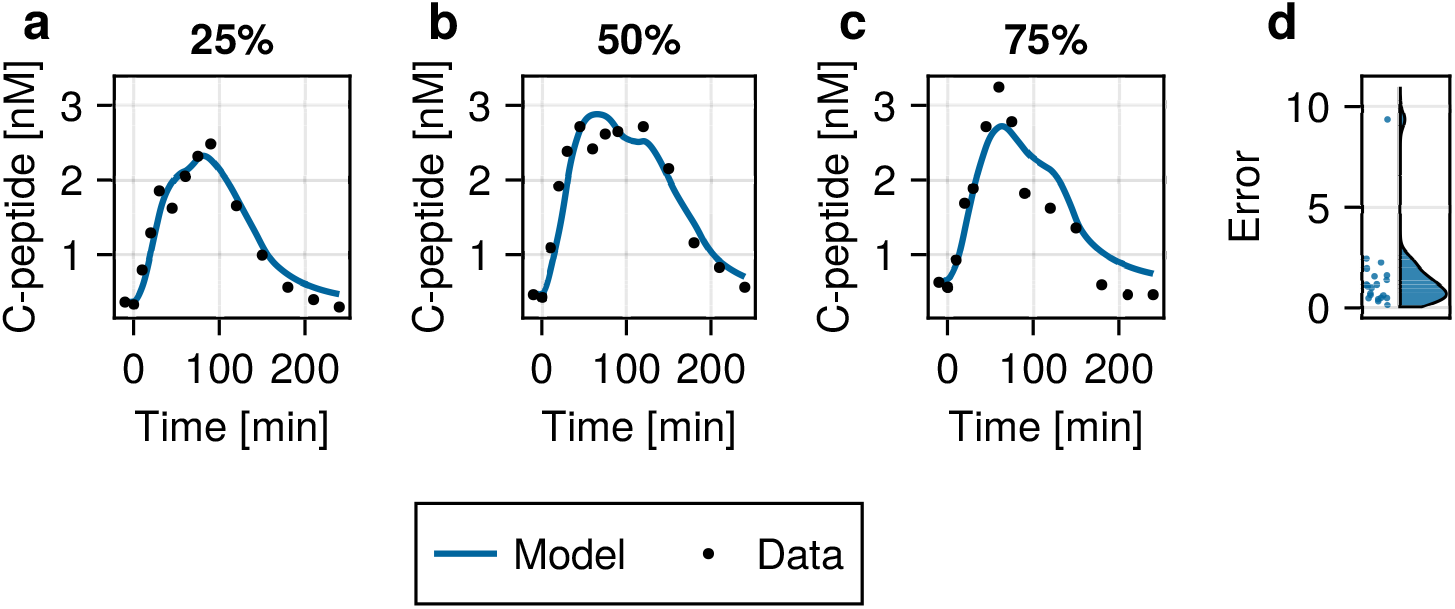
Model fits and errors for the c-peptide model derived using symbolic regression on the Fujita dataset[24]. **a-c**: Model fits for the individuals at the 25^th^, 50^th^ and 75^th^ percentiles of the model error distribution respectively. Measured c-peptide values are indicated with the black circles, the simulated model fits are shown with the dashed line. **d**: Raincloud plot of the model errors for the entire Fujita dataset.

## 3 Discussion

In this work, we introduced conditional universal differential equations (cUDEs) as an extension of the universal differential equation framework that facilitates simultaneous data-driven model discovery and model personalization. We then applied this technique to uncover a novel index of inter-inidividual varition in c-peptide, and by extension, insulin production in a human population with diverse glucose tolerance status.

Our results show that cUDE models can accurately estimate a missing c-peptide production term from the data. More importantly, by accommodating the large interindividual variation in plasma glucose and c-peptide level, the cUDE learns a model that can be generalised across individuals with different glucose tolerance status. In contrast, the classical UDE was unable to generalise from normoglycemic individuals towards impaired glucose tolerance and type 2 diabetes. Investigating individual model fits, the trained cUDE was unable to describe the measurements of c-peptide of a single individual (S4 Fig individual 10) from the test set. However, this individual showed a strong discordance between the measured glucose and c-peptide data with measured plasma glucose only increasing 60 minutes after ingestion of the glucose solution. This unexpected plasma glucose response may potentially be explained by the effect of incretin hormones such as GLP-1 or GIP. These incretin hormones are produced in response to an increase in glucose level in the intestine and activate insulin and c-peptide production [25, 26]. In this study, these hormones are not measured and are a potential additional source of inter-individual variability in c-peptide production. Should time series of incretin hormones become available in the future, the cUDE framework could be reapplied without strong modifications to further learn the role of these incretin hormones in c-peptide production. However, in the current model, where only glucose is provided as the stimulus for c-peptide release, the majority of model fits showed a strong agreement with plasma measurements, suggesting that glucose is the primary driver of c-peptide production. [27]

Furthermore, by constraining the weights and biases of the neural network to be the same for the entire population, the free conditional parameters capture the inter-individual variation which enables the direct comparison between individuals. By comparing the conditional parameters resulting from the c-peptide model with a range of independent measures of metabolic health, we have shown that the conditional parameter strongly correlates with metrics of insulin secretion measured using the hyperglycemic clamp method, the current gold standard measure of insulin production capacity. Furthermore, the lack of a strong correlation with the insulin sensitivity index indicates that the conditional parameter specifically targets the c-peptide and insulin production capacity, and not just a general deterioration in metabolic resilience. The moderate correlation with age may have two causes. Firstly, the conditional parameter has been shown to describe the progressive decline in *β*-cell function, with higher values in people with T2DM. The age distribution was different between each glucose tolerance condition, with the T2DM group being significantly older than the NGT individuals. Secondly, part of this correlation may also originate from the known natural decline of *β*-cell function with ageing. [28]

In order to learn a generalisable model of c-peptide production, a sufficiently heterogeneous dataset is required. Here, we used data from the Ohashi data set consisting of individuals with normal glucose tolerance, imparied gluocse tolerance and T2DM. However, it is not essential to have very large data sets. In S5 Fig we show that using data from 29 individuals did not strongly increase the test error of the cUDE model, provided that the proportion of NGT, IGT and T2DM was maintained. Although the amount of data required also depends on the complexity of the model to be learned, and thereby the neural network size and the amount of inputs and outputs, the cUDE is relatively data-efficient, compared to fully data-driven methods, which typically require thousands of samples. [29, 30]

Furthermore, we show that the interpretability of the cUDE model can also be further increased through the use of symbolic regression. For symbolic regression, we have used a genetic algorithm, which is non-deterministic and may produce variable results upon repeated runs. This can be mitigated by letting the algorithm run through sufficient iterations, which will eventually lead to model convergence. However, this required number of iterations (25,000 in this work) is problem dependent, and for larger problems, more iterations are required, which should be taken into account when applying symbolic regression based on genetic algorithms. In this work, the use of a limited number of allowed operators based on knowledge of previously built ODE models in systems biology greatly reduced the search space, allowing for the discovery of an interpretable model. However, some detail in the dose-response relationship is lost when comparing the analytic expression to the neural network (S8 Fig). Despite the loss of some of the details in the dose response, the derived analytical equation demonstrates generalisability beyond the original dataset, as shown by fitting the derived analytical model to normoglycemic individuals from a previously unseen dataset. Furthermore, we demonstrate that the derived model originally trained on 120 minutes of data can successfully simulate model behaviour over 240 minutes.

Nevertheless, both the Ohashi and Fujita datasets contain only people of Japanese descent. Although previous work has provided evidence for similar *β*-cell responsiveness across all glucose tolerance states [31], it is necessary to further validate the trained model on more diverse populations. In addition, the derived model has only been tested on OGTT responses. Especially considering the effect of amino acids on insulin and c-peptide production [32], the model may not be able to accurately describe responses to more complex meals. However, despite these limitations in the learned c-peptide model, we demonstrate that the cUDE approach outperforms current UDE approaches in learning a generalisable model that incorporates biologically relevant inter-individual variation.

In conclusion, we present cUDEs as an effective extension to the UDE framework that can be used to learn a generalisable representation of missing dynamics from a heterogeneous dataset. The cUDE works under the main assumption that the dynamic system underlying the data is common to all samples, while only a limited set of parameters is necessary to capture the differences between samples. This setup makes the cUDE especially suited to biological challenges, where inter-individual variability is both ubiquitous and often physiologically relevant. Here, we show that the conditional parameter in the cUDE model for c-peptide is interpretable as a physiologically relevant index, capturing the inter-individual variability in c-peptide production as validated by comparison with the hyperglycemic clamp. Although this study demonstrates the application of the cUDE model in a specific application, the cUDE framework is also usable in several other medical disciplines where mathematical models are abundant, such as cardiovascular medicine, neurology, and infectious diseases. The ability of the cUDE model to learn a model that can generalise, capture relevant and interpretable inter-individual variation, and to be trained with limited number of learning examples are key features that demonstrate its potential to support model- and data-driven precision healthcare.

## 4 Methods

### 4.1 Data Collection

#### 4.1.1 Ohashi Dataset

The Ohashi dataset was obtained from Ohashi et al. [33, 34], and originally collected by Okuno et al. [22]. The original study was approved by the ethics committee of the Kobe University Graduate School of Medicine and was registered with the University Hospital Medical Information Network (UMIN000002359). Written informed consent was obtained for all subjects.

As described in [33], 50 subjects with normal glucose tolerance (NGT), 18 subjects with impaired glucose tolerance (IGT), and 53 subjects with type 2 diabetes (T2DM) participated in the study. The characteristics of the subjects for each group are shown in table 1.

**Table 1.** Subject characteristics from the Ohashi dataset [22, 33, 34] after exclusion of subjects with missing data, and the Fujita dataset [24]. For the Ohashi dataset, data is given per glucose tolerance condition.

|  | Ohashi |  |  | Fujita |
| --- | --- | --- | --- | --- |
|  | NGT | IGT | T2DM | - |
| **number of individuals** | 49 | 17 | 51 | 20 |
| **male/female** | 22/27 | 10/7 | 31/20 | 14/6 |
| **age (y)<sup>1</sup>** | 30 ± 8.5 | 41 ± 12 | 55 ± 14 | 29 ± 9 |
| **BMI (kg/m<sup>2</sup>)<sup>1</sup>** | 21 ± 3.4 | 27 ± 6.7 | 26 ± 5.0 | 20.8 ± 2.2 |
| **fasting glucose (mM)<sup>1</sup>** | 4.75 ± 0.43 | 4.90 ± 0.55 | 5.77 ± 0.99 | 5.32 ± 0.44 |
| **2-h glucose (mM)<sup>1</sup>** | 6.14 ± 0.93 | 9.17 ± 0.89 | 14.8 ± 4.24 | 7.27 ± 1.94 |

<sup>1</sup>Data presented as mean ± standard deviation

All subjects underwent a 75-gram oral glucose tolerance test, as well as a consecutive hyperglycemic and hyperinsulinemic-euglycemic clamp test. Both tests were performed on separate mornings after an overnight fast.

In the 75g-OGTT, follwing an overnight fast, blood samples were collected before and 30, 60, 90 and 120 minutes after ingestion of the glucose solution. Plasma glucose and serum insulin and c-peptide concentrations were measured in each sample.

Hyperglycemic clamp and hyperinsulinemic-euglycemic clamp tests were performed consecutively. The hyperglycemic clamp began with an intravenous infusion of a glucose bolus of 9622 mg m^*−*2^ within 15 minutes, followed by a variable dose of glucose to keep plasma glucose levels at 200 mg dL^*−*1^ for 90 minutes. Blood samples were collected before and at 5, 10, 15, 60, 75, and 90 minutes after glucose infusion. In each blood sample, plasma glucose and serum insulin and c-peptide were measured. The hyperinsulinemic-euglycemic clamp test was then performed by intravenous infusion of regular human insulin at 1.46 mU kg^*−*1^ min^*−*1^ to obtain a serum insulin concentration of 600 pmol L^*−*1^. Plasma glucose concentration was kept at 90 mg dL^*−*1^ by variable glucose infusion for 120 minutes. [22]

Insulin secretion indices were defined as the incremental area under the insulin concentration curve during the hyperglycemic clamp:

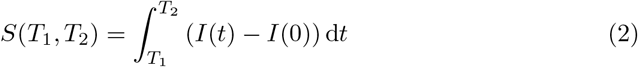

Where the insulin secretion during the first-phase is defined as *S*(0, 10), during the second phase as *S*(10, 90) and the total insulin secretion as *S*(0, 90). The insulin sensitivity index (ISI) is calculated from the hyperinsulinemic-euglycemic clamp by dividing the mean measured glucose infusion rate during the last 30 minutes of the test by the product of plasma glucose and serum insulin levels at the end of the clamp (*t* = 120).

#### 4.1.2 Fujita Dataset

The Fujita dataset was obtained from Fujita et al. [24]. Written informed consent was obtained for all subjects.

As described in [24], 20 subjects with normal glucose tolerance (NGT) participated in the study. Subject characteristics for each group are shown in table 1. All subjects underwent a 75g-oral glucose tolerance test (OGTT) in the morning after an overnight fast. Fasting blood samples were drawn twice before oral ingestion of glucose. Blood samples were obtained at 10, 20, 30, 45, 60, 75, 90, 120, 150, 180, 210, 240 min after ingestion. Subjects remained at rest throughout the test. Blood samples were rapidly centrifuged. [24]

### 4.2 Data Preprocessing

Measurements of four subjects with missing values in the OGTT experiment were excluded from further analysis. The values reported in table 1 are calculated on the data after exclusion. Unit conversions were performed to convert glucose from mg dL^*−*1^ to mM and c-peptide from ng mL^*−*1^ to nM. For the data from Fujita et al. [24], no measurements were dropped and the same unit conversions were applied, as with the data from Ohashi et al.

### 4.3 Implementation Details

#### 4.3.1 Differential Equation Model of C-Peptide

The van Cauter model was used to describe the concentrations of c-peptide in the plasma and interstitial compartment, [21] (figure 2a). The original model, used to describe intravenously administered c-peptide, was extended to include endogenous production of c-peptide by the pancreas. The model equations for both compartments are given by:

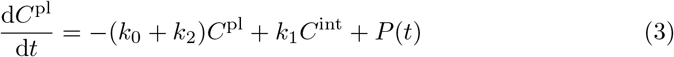

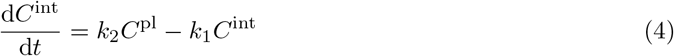

Where *C*^pl^ represents the concentration of c-peptide in the plasma compartment and *C*^int^ is the concentration of C-peptide in the interstitial compartment. Kinetic parameters *k*_0_-*k*_2_ were calculated for each individual based on age, using equations provided by van Cauter et al. [21], which are given as:

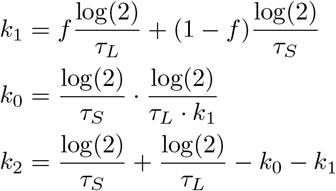

For which the parameter values (*f, τ*_*S*_, and *τ*_*L*_) are given in table 2 for the NGT, IGT, and T2DM groups.

**Table 2.** Parameter values for computing the kinetic parameters for the van Cauter c-peptide model for the NGT, IGT and T2DM groups.

| Parameter (unit) | Value (NGT, IGT) | Value (T2DM) |
| --- | --- | --- |
| $f$ (-) | 0.76 | 0.78 |
| $\tau_S$ (min) | 4.95 | 4.52 |
| $\tau_L$ (min) | $29.2 + 0.14 \cdot \text{Age}$ | |

#### 4.3.2 Neural Network Component

The production of c-peptide *P*(*t*) was modelled using a densely connected neural network with two inputs; the first was given by the difference in plasma glucose at time *t* compared to fasting values (*G*_*i*_(*t*) = *G*^pl^(*t*) *− G*^pl^(0)) and the second was a learnable parameter *β*_*i*_ representing the inter-individual variability (fig. 2b). Plasma glucose values are obtained directly from the measured data using a forcing function. For timepoints in between measurements, glucose values are linearly interpolated.

The neural network contained two hidden layers each consisting of 6 nodes making use of tanh-activation functions, and an output layer of size 1, with a softplus activation function, resulting in 67 trainable weights.

The neural network architecture was selected through a grid search. Different architectures were obtained by varying the depth of the model between 1 to 2 layers, with layer sizes of 3 to 6 nodes, and 3 layers with layer sizes of 3 and 4 nodes. All models were trained on 70% of the train set and evaluated in the remaining 30%. The model that gave the lowest error for most individuals was selected. In case of a tie, the model with the lowest median error on all individuals was selected.

#### 4.3.3 Initial Conditions

For simulation, the whole system was assumed to be in steady state at *t* = 0, as subjects were fasting prior to the oral glucose tolerance test. The initial condition for plasma c-peptide (*C*^pl^) was set to the measured fasting value at *t* = 0. For interstitial c-peptide, the initial condition was calculated using the steady-state assumption to be

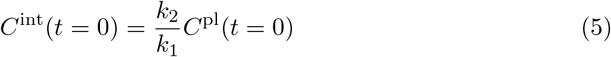

Furthermore, to ensure the plasma c-peptide compartment was in steady-state at *t* = 0, the production term *P*(*t*) including the neural network *N* (*G, β*_*i*_), describing c-peptide production was formulated as

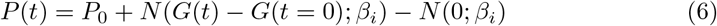

Resulting in a production value at *t* = 0 of *P* (*t* = 0) = *P*_0_, where *P*_0_ was set as

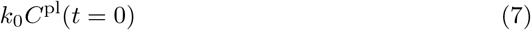

### 4.4 Parameter Estimation

The neural network parameters were estimated on a randomly selected training subset containing 70% of the total samples, stratified according to the glucose tolerance condition. This training set was further divided into a true train set of 70% and a validation set of 30% of samples. This resulted in a true training set containing 49% of the entire dataset (*n* = 57), and a validation set containing 21% of the entire dataset (*n* = 25). Parameters were estimated on the true train set using the following loss function:

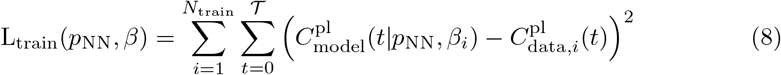

Where *p*_NN_ are the parameters of the neural network, *β* represents the vector of all conditional parameters for each individual *i* out of the total of *N*_train_ individuals. Furthermore, *T* = *{*0, 30, 60, 90, 120*}* represents the set of timepoints contained in the data.

The parameter estimation for the universal differential equation models was then performed by sampling 25,000 initial candidate parameter sets and optimising the 25 candidate parameter sets that yielded the smallest initial objective function values. Subsequent optimisation was performed using a two-stage optimiser, starting with Adam [35] for 1,000 iterations with a learning rate of 10^*−*2^. Starting from the endpoint of Adam, the LBFGS optimiser was used for a maximum of 1,000 iterations or until convergence. Subsequently, for all trained 25 models, the neural network parameters were fixed and the conditional parameters were estimated on the validation set using the LBFGS optimiser, with the following loss function:

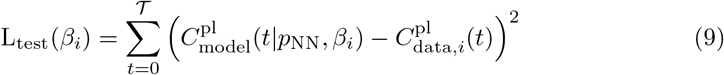

The model that resulted in the lowest loss function value in the majority of individuals in the validation set was then selected as the best performing model.

For the remaining 30% of the complete data set (*n* = 35), labelled the test set, only the conditional parameter was estimated. In this test subset, the parameters for each individual were estimated separately using the LBFGS algorithm only, and using the loss function in equation 9.

### 4.5 Symbolic Regression

For symbolic regression, initially 900 unique samples of the neural network output were created through combinations of 30 values for the conditional parameter *β* and incremental glucose values. Incremental glucose values were capped at zero, to reduce the complexity of the problem. Symbolic regression was then performed using the PySR package [23] using the settings listed in table 3.

**Table 3.**
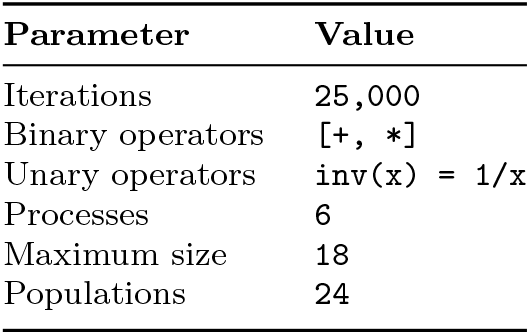
Settings for the symbolic regression algorithm from PySR.

From the resulting equations, the top equation was selected using the ‘best’ option from the PySR package. This first selects all expressions with a loss smaller than at least 1.5 times the loss of the most accurate model. From these expressions, the model equation with the highest score is selected, defined as the negated derivative of the loss with respect to complexity. [23]

The resulting equation was then simplified by amalgamating constants into a single learnable parameter. As the incremental glucose values used to train the symbolic equation were capped at zero, production was set at a value of zero when *G*^pl^(*t*) *< G*^pl^(0).

#### 4.5.1 Programming

Both the ordinary differential equation models, as well as the universal differential equation models used in this research were implemented in the Julia programming language, using the ‘OrdinaryDiffEq.jl’ package [36].

## Declarations

## Code and data availability

All code and data used to produce the results and analyses can be found in the GitHub repository linked to this publication: https://github.com/Computational-Biology-TUe/conditional-ude

## Author contributions

**Max de Rooij:** Conceptualisation, Methodology, Software, Investigation, Formal analysis, Writing - original draft, Visualisation. **Natal A.W. van Riel:** Conceptualisation, Resources, Writing - Review & Editing, Supervision. **Shauna D. O’Donovan:** Conceptualisation, Resources, Writing - Review & Editing, Supervision, Project administration, Funding acquisition.

## Supplementary information

**S1 Fig.**
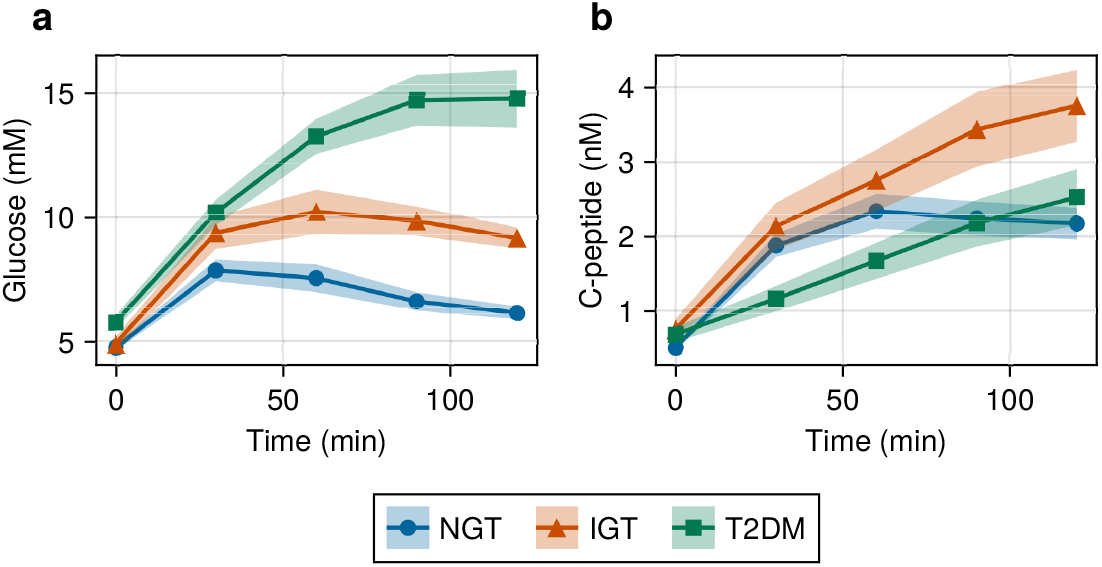
Time-series data of measured plasma glucose and c-peptide from the Ohashi et al. dataset [33, 34]. **(a)** Time-series data of plasma glucose after an OGTT, separated by glucose tolerance status. **(b)** Time-series data of plasma c-peptide after an OGTT, separated by glucose tolerance status. Solid lines and markers show the mean values for each subgroup and shaded regions indicate 1.96 times the sub-population standard error. Blue (circles) normal glucose tolerance (NGT), orange (triangles) are impaired glucose tolerance (IGT), and green (squares) represent the type 2 diabetes mellitus (T2DM) group.

**S2 Fig.**
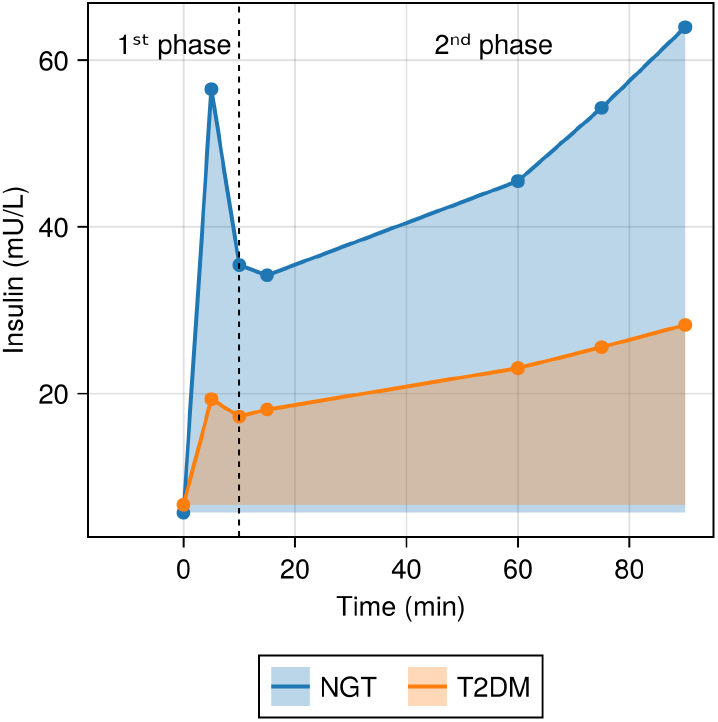
Illustration of the hyperglycemic clamp data. During the experiment, hyperglycemia is induced by a glucose infusion, raising plasma glucose to 6.9 mM above basal levels, and maintaining the hyperglycemic plateau by dynamically adjusting the glucose infusion rate. The resulting insulin concentrations provide a measure of the pancreatic *β*-cell capacity. The profile is divided into a first phase (0-10 min) and a second phase (10-90 min). For each phase, the area under the insulin curve can be computed to yield a measure of the first- and second-phase insulin response respectively. In this figure, the difference between normal glucose tolerance (NGT) and type 2 diabetes (T2DM) is illustrated.

**S3 Fig.**
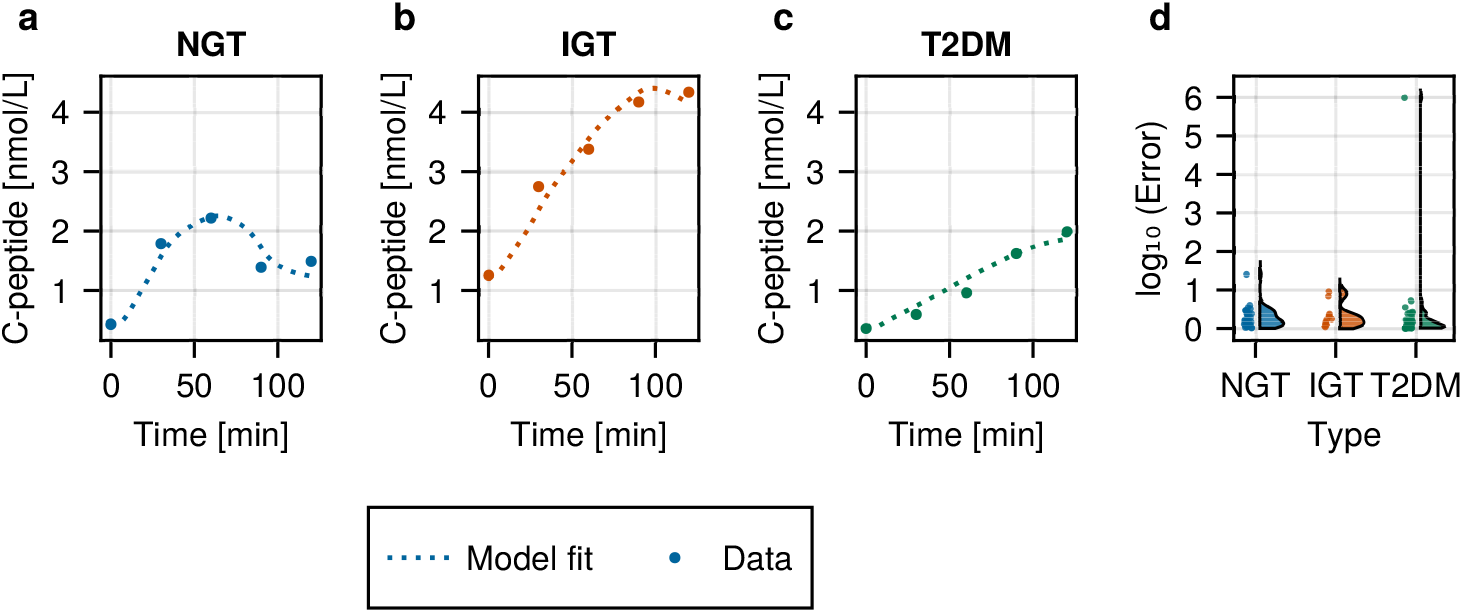
Model fits of the conditional UDE model on the train data. **a-c**: Model fit of the individuals in the train set with the median error value within each glucose tolerance status group. **d**: Distribution of error values for model fits for all subjects in the train subset, separated by glucose tolerance status group.

**S4 Fig.**
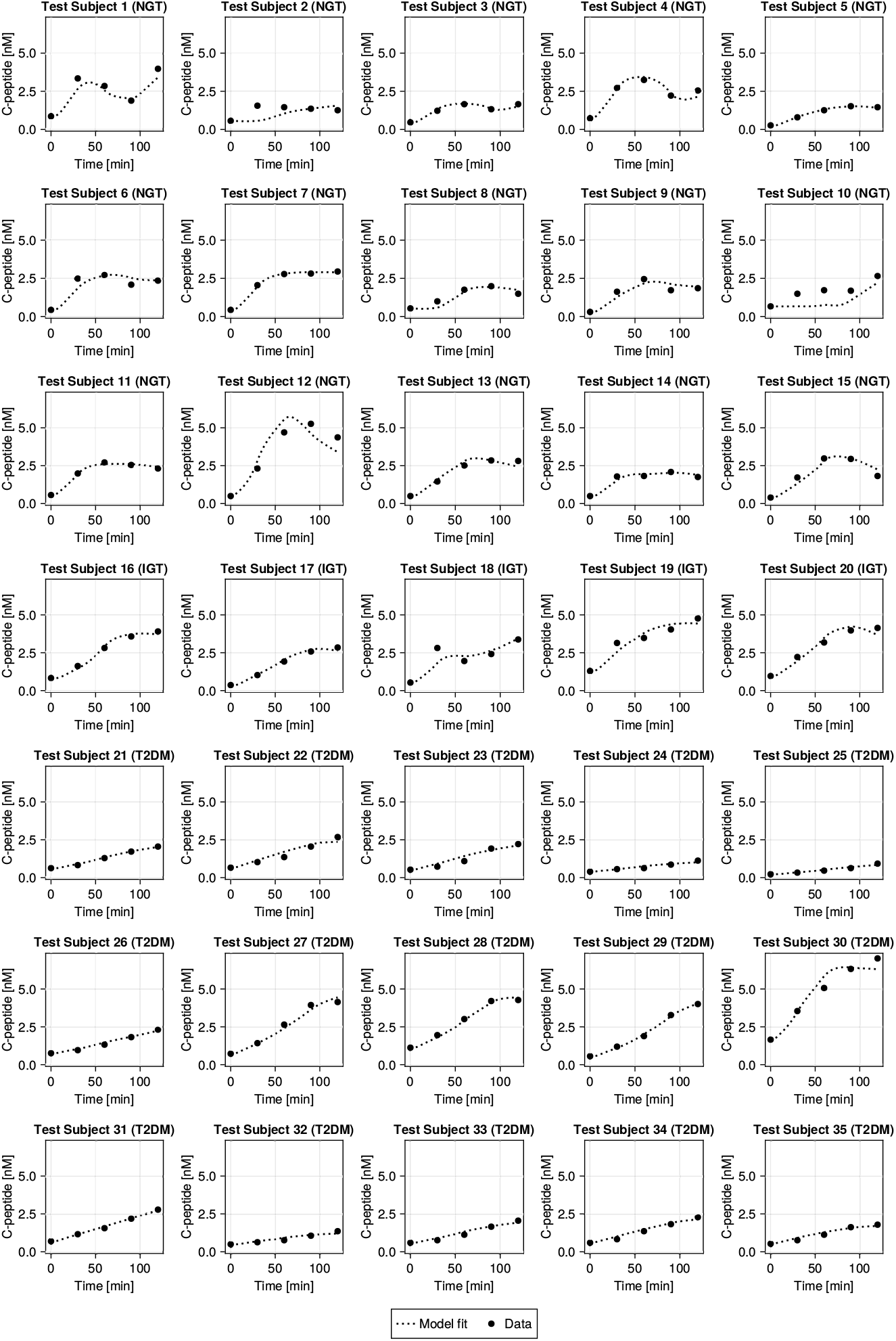
All model fits of the cUDE model on the test data. The dashed line indicates the model fit and the dots indicate the c-peptide measurements. The subgroup (NGT, IGT, or T2DM) is included after each subject number in the title of each subplot.

**S5 Fig.**
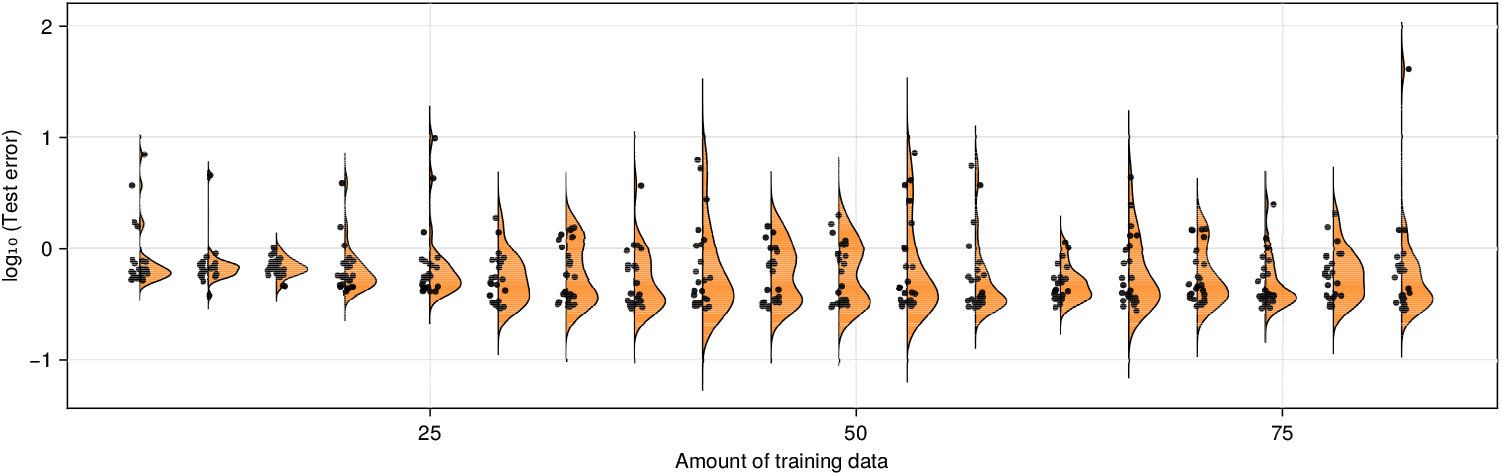
Test error for models trained on a subset of the training data. The cUDE model was trained on a fraction of the training data ranging from 10 to 100%, stratified by glucose tolerance status. The resulting models were evaluated on the test set and the test error was computed. The solid line indicates the mean test error, and the shaded area represents the mean *±*1.96 *·* Standard error.

**S6 Fig.**
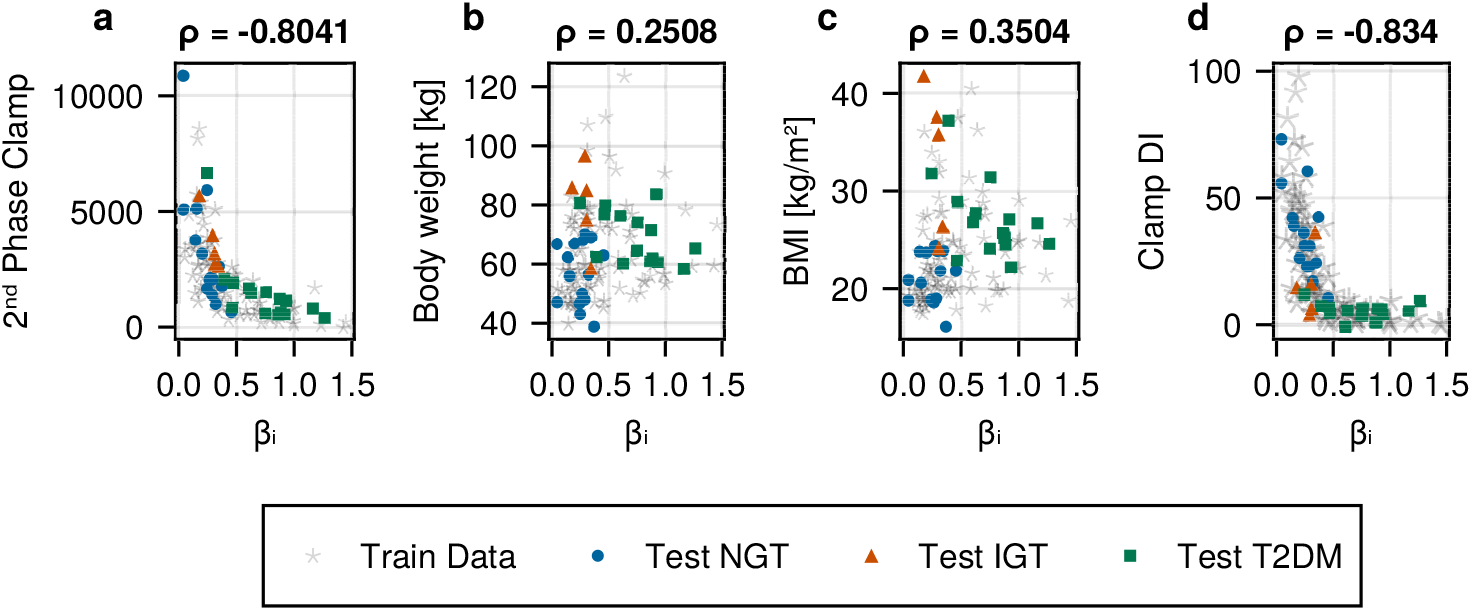
Correlation of cUDE conditional parameter with metrics of insulin production capacity and physiological measurements. Correlation of *β*_*i*_ with (a) second-phase clamp insulin auc, (b) body weight, (c) BMI, and (d) Clamp disposition index. The grey circles indicate the train data. The blue circles represent the normal glucose tolerance individuals, the orange triangles are the impaired glucose tolerance individuals, and the green squares represent the individuals with type 2 diabetes mellitus.

### S7. Simplification of the Symbolic Regression Equation The equation resulting directly from the symbolic regression algorithm was formulated as

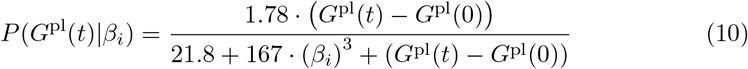

As *β*_*i*_ is a trainable parameter, we can replace it with *k*_*M*_ defined as

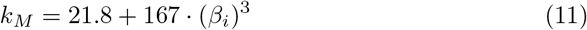

Setting the production to zero for *G*^pl^(*t*) *< G*^pl^(0) then results in equation 1.

**S8 Fig.**
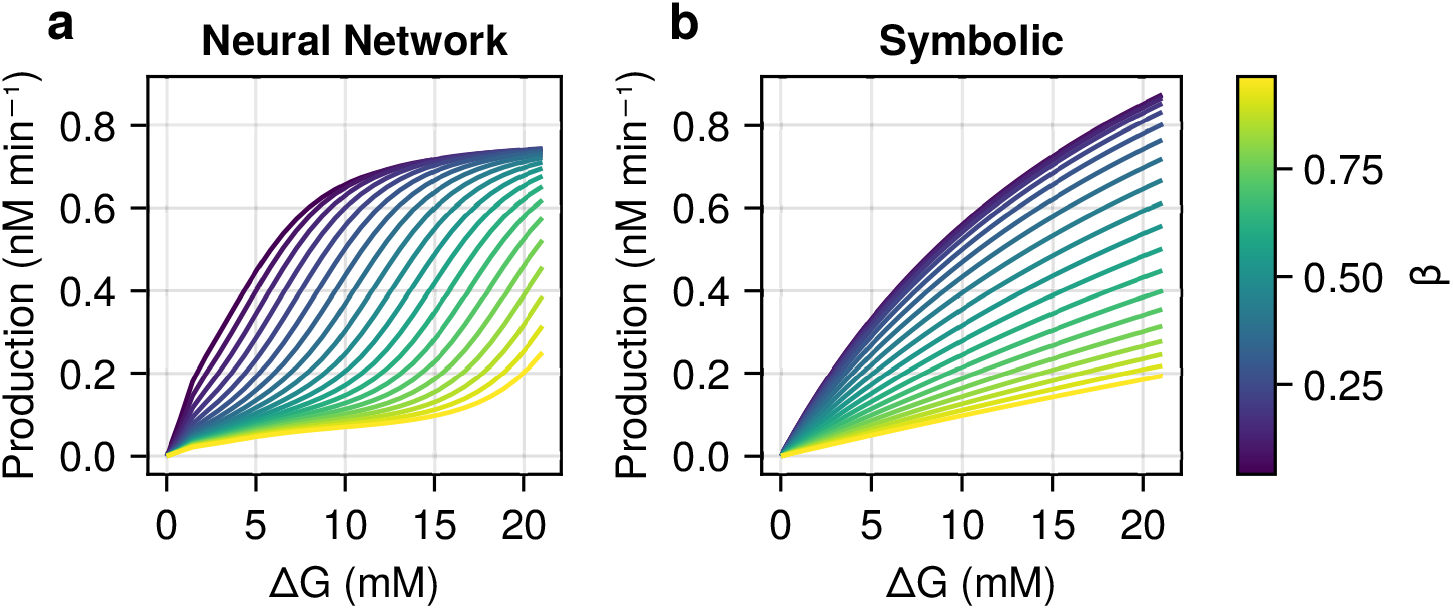
Dose-response curves for varying *β* and glucose values for the a: neural network and b: the derived analytic expression. The values for *β* are aligned by setting *k*_*M*_ in the analytical expression according to equation 11.

## References

[1] Nielsen, J. Systems biology of metabolism: A driver for developing personalized and precision medicine. Cell Metabolism 25 (2017).

[2] Wysham, C. & Shubrook, J. Beta-cell failure in type 2 diabetes: mechanisms, markers, and clinical implications. Postgraduate Medicine 132 (2020).

[3] Jones, A. G. et al. Markers of β-cell failure predict poor glycemic response to glp-1 receptor agonist therapy in type 2 diabetes. Diabetes Care 39 (2016).

[4] Thong, K. Y. et al. The association between postprandial urinary c-peptide creatinine ratio and the treatment response to liraglutide: A multi-centre observational study. Diabetic Medicine 31 (2014).

[5] Ashley, E. A. Towards precision medicine. Nature Reviews Genetics 17 (2016).

[6] Coral, D. E. et al. Subclassification of obesity for precision prediction of cardiometabolic diseases. Nature Medicine (2024). URL 10.1038/s41591-024-03299-7.

[7] Berger, M. F. & Mardis, E. R. The emerging clinical relevance of genomics in cancer medicine. Nature Reviews Clinical Oncology 15 (2018).

[8] Finotello, F. & Eduati, F. Multi-omics profiling of the tumor microenvironment: Paving the way to precision immuno-oncology. Frontiers in Oncology 8 (2018). URL https://www.frontiersin.org/article/10.3389/fonc.2018.00430/full.

[9] Wang, H. et al. Deep learning in systems medicine. Briefings in Bioinformatics 22 (2021).

[10] Ching, T. et al. Opportunities and obstacles for deep learning in biology and medicine. Journal of the Royal Society Interface 15 (2018).

[11] Sapoval, N. et al. Current progress and open challenges for applying deep learning across the biosciences. Nature Communications 2022 13:1 13, 1–12 (2022). URL https://www.nature.com/articles/s41467-022-29268-7.

[12] Rudin, C. Stop explaining black box machine learning models for high stakes decisions and use interpretable models instead. Nature Machine Intelligence 1 (2019).

[13] Lauzeral, N. et al. A model order reduction approach to create patient-specific mechanical models of human liver in computational medicine applications. Comput. Methods Programs Biomed. 170, 95–106 (2019).

[14] Kuepfer, L. & Schuppert, A. Systems medicine in pharmaceutical research and development. Methods Mol. Biol. 1386, 87–104 (2016).

[15] O’Donovan, S. D. et al. Quantifying the effect of nutritional interventions on metabolic resilience using personalized computational models. iScience 27, 109362 (2024). URL https://linkinghub.elsevier.com/retrieve/pii/S2589004224005832.

[16] Erdős, B. et al. Quantifying postprandial glucose responses using a hybrid modeling approach: Combining mechanistic and data-driven models in the maastricht study. PLOS ONE 18, e0285820 (2023). URL https://journals.plos.org/plosone/article?id=10.1371/journal.pone.0285820.

[17] Raissi, M., Perdikaris, P. & Karniadakis, G. E. Physics-informed neural networks: A deep learning framework for solving forward and inverse problems involving nonlinear partial differential equations. Journal of Computational Physics 378, 686–707 (2019).

[18] Rackauckas, C. et al. Universal differential equations for scientific machine learning. ArXiv (2021).

[19] de Rooij, M., Erdős, B., van Riel, N. A. W. & O’donovan, S. D. Physiology-informed regularization enables training of universal differential equation systems for biological applications. bioRxiv 2024.05.28.596164 (2024). URL https://www.biorxiv.org/content/10.1101/2024.05.28.596164v1https://www.biorxiv.org/content/10.1101/2024.05.28.596164v1.abstract.

[20] Philipps, M., Körner, A., Vanhoefer, J., Pathirana, D. & Hasenauer, J. Non-negative universal differential equations with applications in systems biology. ArXiv (2024). URL https://arxiv.org/abs/2406.14246v1.

[21] van Cauter, E., Mestrez, F., Sturis, J. & Polonsky, K. S. Estimation of insulin secretion rates from c-peptide levels. comparison of individual and standard kinetic parameters for c-peptide clearance. Diabetes 41, 368–377 (1992). URL https://pubmed.ncbi.nlm.nih.gov/1551497/.

[22] Okuno, Y. et al. Postprandial serum c-peptide to plasma glucose concentration ratio correlates with oral glucose tolerance test- and glucose clamp-based disposition indexes. Metabolism: Clinical and Experimental 62, 1470–1476 (2013). URL http://www.metabolismjournal.com/article/S0026049513001753/fulltexthttp://www.metabolismjournal.com/article/S0026049513001753/abstracthttps://www.metabolismjournal.com/article/S0026-0495(13)00175-3/abstract.

[23] Cranmer, M. Interpretable machine learning for science with pysr and symboli-cregression.jl. ArXiv (2023). URL https://arxiv.org/abs/2305.01582v3.

[24] Fujita, S. et al. Four features of temporal patterns characterize similarity among individuals and molecules by glucose ingestion in humans. npj Systems Biology and Applications 2022 8:1 8, 1–16 (2022). URL https://www.nature.com/articles/s41540-022-00213-0.

[25] Holst, J. J., Gasbjerg, L. S. & Rosenkilde, M. M. The role of incretins on insulin function and glucose homeostasis. Endocrinology (United States) 162 (2021).

[26] Pasveer, Y. M. et al. Does GLP-1 cause post-bariatric hypoglycemia: ‘Computer says no’. Computer Methods and Programs in Biomedicine 257, 108424 (2024). URL https://www.sciencedirect.com/science/article/pii/S0169260724004176.

[27] Henquin, J. C. Triggering and amplifying pathways of regulation of insulin secretion by glucose. Diabetes 49, 1751–1760 (2000). URL 10.2337/diabetes.49.11.1751.

[28] Aguayo-Mazzucato, C. Functional changes in beta cells during ageing and senescence. Diabetologia 63 (2020).

[29] Zeevi, D. et al. Personalized nutrition by prediction of glycemic responses. Cell (2015).

[30] Lutsker, G. et al. From glucose patterns to health outcomes: A generalizable foundation model for continuous glucose monitor data analysis. ArXiv (2024).

[31] Møller, J. B. et al. Ethnic Differences in Insulin Sensitivity, β-Cell Function, and Hepatic Extraction Between Japanese and Caucasians: A Minimal Model Analysis. The Journal of Clinical Endocrinology & Metabolism 99, 4273–4280 (2014). URL 10.1210/jc.2014-1724.

[32] van Sloun, B. et al. The Impact of Amino Acids on Postprandial Glucose and Insulin Kinetics in Humans: A Quantitative Overview. Nutrients 12, 3211 (2020). URL https://www.mdpi.com/2072-6643/12/10/3211. Number: 10 Publisher: Multidisciplinary Digital Publishing Institute.

[33] Ohashi, K. et al. Increase in hepatic and decrease in peripheral insulin clearance characterize abnormal temporal patterns of serum insulin in diabetic subjects. npj Systems Biology and Applications 2018 4:1 4, 1–12 (2018). URL https://www.nature.com/articles/s41540-018-0051-6.

[34] Ohashi, K. et al. Glucose homeostatic law: Insulin clearance predicts the progression of glucose intolerance in humans. PLOS ONE 10, e0143880 (2015). URL https://journals.plos.org/plosone/article?id=10.1371/journal.pone.0143880.

[35] Kingma, D. P. & Ba, J. L. Adam: A method for stochastic optimization. 3rd International Conference on Learning Representations, ICLR 2015 - Conference Track Proceedings (2014). URL https://arxiv.org/abs/1412.6980v9.

[36] Rackauckas, C. & Nie, Q. Differentialequations.jl – a performant and feature-rich ecosystem for solving differential equations in julia. Journal of Open Research Software 5, 15 (2017). URL https://openresearchsoftware.metajnl.com/articles/10.5334/jors.151.

